# Unravelling bird nest arthropod community structure using metabarcoding

**DOI:** 10.1101/2023.03.09.531929

**Authors:** Valerie Levesque-Beaudin, Dirk Steinke, Mieke Böcker, Bettina Thalinger

**Author notes:** Corresponding author: Bettina Thalinger.

## Abstract

Bird nests are fascinating microcosms harboring a wide range of arthropods parasitizing the nesting birds or feeding on prey remains, feces, and the nest material. Studies of these communities have been entirely based on emergence traps which collect live organisms out of the nests. The analysis of nest contents and environmental DNA (eDNA) via metabarcoding could expand our knowledge and identify prey, exuviae, and other animal remains in bird nests.

Here, we investigated the potential of arthropod remains, nest dust, and feathers to better describe taxonomic diversity accumulated in 20 bird nests collected in Guelph (Canada). We used subsampling strategies and tested two extraction approaches to investigate the distribution of DNA in nests, account for low-quality DNA, and the presence of inhibitory substances.

In total, 103 taxa were detected via metabarcoding. Arthropod remains delivered the highest number of taxa (n=67), followed by nest dust (n=29). Extractions with the PowerSoil kit outperformed DNeasy extractions coupled with PowerClean Pro inhibitor removal. The subsamples of the same nest showed 5.5% and 47.1% taxonomic overlap for arthropod remains and PowerSoil extracted nest dust, respectively, indicating a heterogeneous eDNA distribution in nests. Most detected species were either feeding in the nest, i.e., herbivorous / predatory, or bird food. We also detected molecular traces of 25 bird species, whose feathers were likely used as nest material.

Consequently, the metabarcoding of bird nest materials provides a more complete picture of nest communities, which can enable future studies on functional diversity and better comparisons between nesting species.

## Introduction

A bird nest is a fascinating microcosm. To the naked eye, it appears to consist mainly of plant matter, branches, feathers, and sometimes mud, but it is also home to a small cosmos of arthropods. Upon closer inspection, the nest becomes alive with crawling larvae and adult critters holding a wide range of species representing many orders and functional groups (Hicks 1959, 1962, 1971; Woodroffe 1953). However, the study of this particular diversity is not without challenges. As researchers aim to keep disturbance of the birds at minimum, access to the nest is usually limited to the time after fledging, reducing the amount of information available. In addition, many of the invertebrate inhabitants are at immature life stages, making it difficult to gain identification below family level (Pfenninger et al., 2007; Sinclair & Gresens, 2008). Therefore, earlier studies reared arthropods to adulthood for species identification by using rearing chambers or by transferring the entire nest into an emergence trap (Gilbert 2011; Levesque-Beaudin *et al*. 2020). Unfortunately, this approach is limited to taxa still present in the nest at the end of the breeding season.

An alternative could be an approach such as DNA barcoding (Hebert *et al*. 2003) utilizing all animal remains as well as nest dust to provide reliable species level identifications. Especially the use of DNA barcoding for bulk samples, i.e. metabarcoding, promises the rapid determination of the species composition of entire communities (Ritter *et al*. 2019; Steinke *et al*. 2021, 2022) even from remains and fragments (Elbrecht *et al*. 2021b; Ruppert *et al*. 2019). As metabarcoding has never been used for investigating the invertebrate community in a bird nest, there are some important considerations. First, which material should be sampled and what laboratory protocols should be used? Birds use a wide range of nest building materials (e.g., twigs, sticks, branches, mud, pebbles, grass, leaf litter). Some of these, particularly plant material containing humic acids, could induce PCR inhibition (Opel *et al*. 2010; Sutlović *et al*. 2005), others will only contain DNA of low quality (Ruppert *et al*. 2019). Both issues can be addressed by adjusting lab protocols using commercial inhibitor removal kits and different DNA extraction protocols respectively. Secondly, nests are fairly large which raises the question which subsampling strategy should be used? Subsamples could be arthropod remains picked by hand, dust retrieved through sieving, or even feathers or bird feces. Arthropod remains such as body parts or exuviae can considerably vary in size and DNA content, which can be the source of bias as it has been repeatedly shown that metabarcoding is susceptible to biomass variation (e.g., Elbrecht *et al*. 2017a; Elbrecht & Leese 2015). Dust (or debris) contains animal environmental DNA (eDNA) in the form of smaller body parts, minute animals, which are mixed with plant matter, and other fine particles of solid matter (Foster et al., 2023; Lennartz et al., 2021). Similar samples have been successfully analysed using metabarcoding (Madden *et al*. 2016), making them a potentially suitable option. Many species of birds incorporate feathers into their nests. Feathers are known to host several groups of parasitic arthropods (Fryderyk & Izdebska 2009) but can also serve as direct DNA source to confirm the identity of the nesting species. Metabarcoding of bird feces has been used to determine the diet of species and disentangle bird-insect food webs (e.g., Rytkönen *et al*. 2019; Volpe *et al*. 2022). However, given that nest sanitation is a widespread behaviour among birds (Guigueno & Sealy 2012), it is rather unlikely that sufficient quantities of feces are present in a nest at the end of the breeding season for a thorough dietary analysis.

Given these considerations, our aims for this study were to: 1) compare an emergence trap approach with DNA metabarcoding to determine, if the latter can identify a broader range of taxa; 2) compare different types of subsampled material (arthropod remains, eDNA contained in dust, feathers) to determine an efficient and sensitive strategy to investigate species communities in bird nests in future large-scale studies; 3) test two DNA extraction methods and an inhibitor removal protocol for their capability to produce high quality results from bird nest dust samples.

## Materials and Methods

### Bird nest collection and arthropod emergence

In total, we collected 20 nests from six different bird species (Table 1). Of those, 16 were in nest boxes and four were found over ledges in open areas (American Robin: *Turdus migratorius* Linnaeus and Eastern Phoebe: *Sayornis phoebe* (Latham)). Nests were collected between June 12 and June 28, 2019, as well as on October 5, 2019 at the University of Guelph Arboretum (Guelph, Ontario, Canada, 43.541299, −80.214540) after the fledglings had left or at the end of the breeding period, respectively. Additionally, three abandoned nests were collected in Guelph in July 2018, (Table 1). Nests were individually stored in plastic bags and transported back to the lab.

**Table 1.**
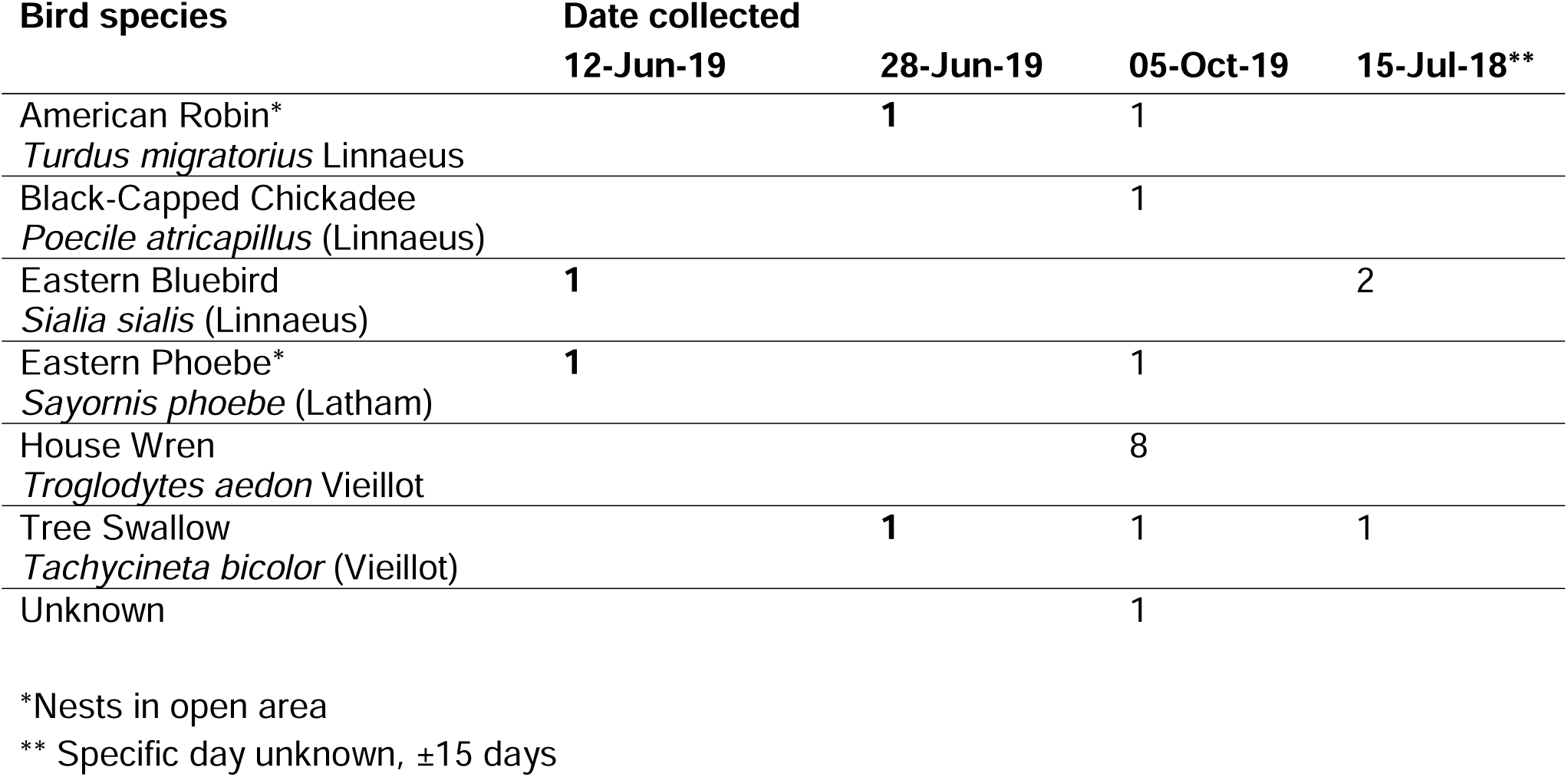
Nests for each bird species by date of collection. Numbers in bold indicate nests processed in emergence traps.

Nests collected during June, were transferred into an emergence trap combined with a Berlese-Tullgren funnel (Fig. 1) and left there for a period of 2-3 weeks to allow emergence of arthropods. Contrary to most Berlese-Tullgren funnel set ups, no extra light source was used in order to slow the arthropods’ downward movement and to allow for more time for emergence. Arthropods were collected both in the Berlese-Tullgren funnel and in the collecting bottle of the emergence trap. They were transferred into 95% ethanol for preservation. Both samples were combined per nest for subsequent DNA barcoding. After the emergence period, each nest was transferred into a fresh plastic bag and frozen at −20°C until further processing in the laboratory.

**Figure 1.**
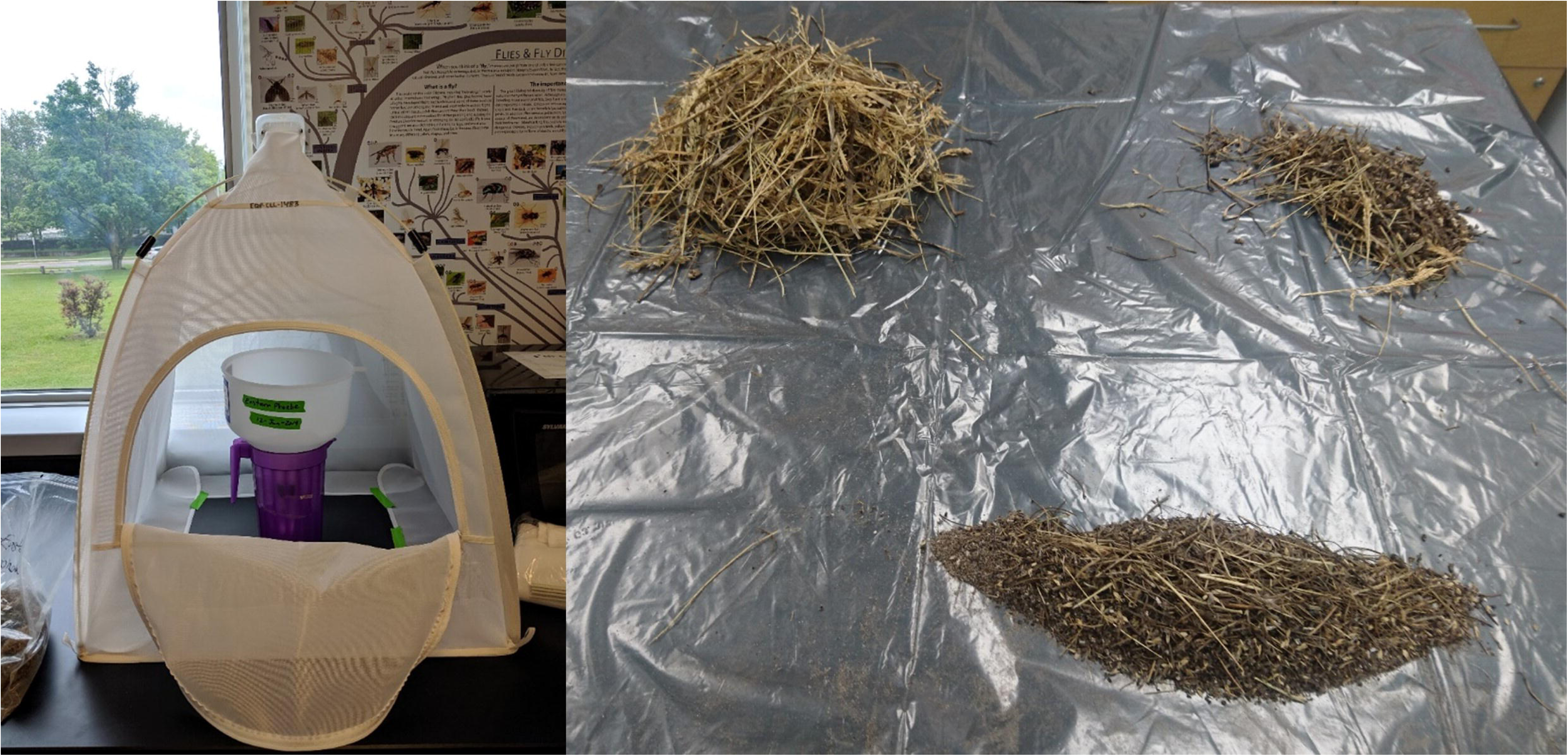
Emergence trap with Berlese-Tullgren funnel (left picture) and nest separated into components >5 mm, >2 mm and <2 mm via consecutive sieving. Individually removed arthropod remains are not displayed.

### DNA barcoding of emerged arthropods

All arthropods were identified to the lowest taxonomic level possible. Five specimens or less per morphospecies were selected for DNA barcoding. One or two legs were removed from each specimen for DNA extraction. Lab work followed standardized protocols for DNA extraction, barcode amplification and sequencing (deWaard et al., 2008). DNA was extracted using a glass-fiber column-based protocol (Ivanova et al., 2006). The primer cocktail C_LepFolF and C_LepFolR (Hernandez Triana et al., 2014) was used to amplify a 658 bp fragment of the COI gene. The PCR thermal regime consisted of an initial denaturation at 94°C for 1 min; five cycles at 94°C for 1 min, 45°C for 1.5 min and 72°C for 1.5 min; 35 cycles of 94°C for 1 min, 50°C for 1.5 min and 72°C for 1 min followed by a final cycle at 72°C for 5 min. PCR amplicons were visualized on a 1.2% agarose gel E-Gel® (Invitrogen) and then diluted 1:10 with sterile water. Amplicons (2–5 μL) were bidirectionally sequenced using sequencing primers M13F or M13R (Messing, 1983) and the BigDye® Terminator v.3.1 Cycle Sequencing Kit (Applied Biosystems, Inc.) on an ABI 3730xl capillary sequencer at the Canadian Centre for DNA Barcoding (CCDB, https://ccdb.ca/). All records were uploaded to the Barcode of Life Data System (BOLD, http://www.boldsystems.org/) and are publicly available through the dataset DS-BRDNT (doi: dx.doi.org/10.5883/DS-BRDNT) and GenBank (accession: OP587213-OP587227; OP599355-OP599544).

### Nest dissection and sieving

Nests were thawed and then sieved twice using 5 mm and 2 mm sieves respectively (Fig. 1). Nest components larger than 5 mm were manually sorted for arthropod remains; these were collectively stored in 5 ml or 50 ml tubes. Fragments too large to fit through the 2 mm sieve were processed alike. Nest components passing the 2 mm sieve were scooped and placed in 500 ml wide neck Nalgene bottles. Additionally, all feathers found in a nest sample were transferred into a single 5 ml or 50 ml tube. This led to a total of 19 bottles (500 ml), 27 larger tubes (50 ml) and 16 smaller tubes (5 ml) which were stored at −20°C until further processing in a low-DNA environment (Fig. 2). All work was done on DNA-free surfaces (cleaned with 1.5% bleach and 70% ethanol) and by using DNA-free gloves, which were changed between nests. Sieves and forceps were cleaned with 5% bleach, rinsed with water, and dried after work on a nest.

**Figure 2.**
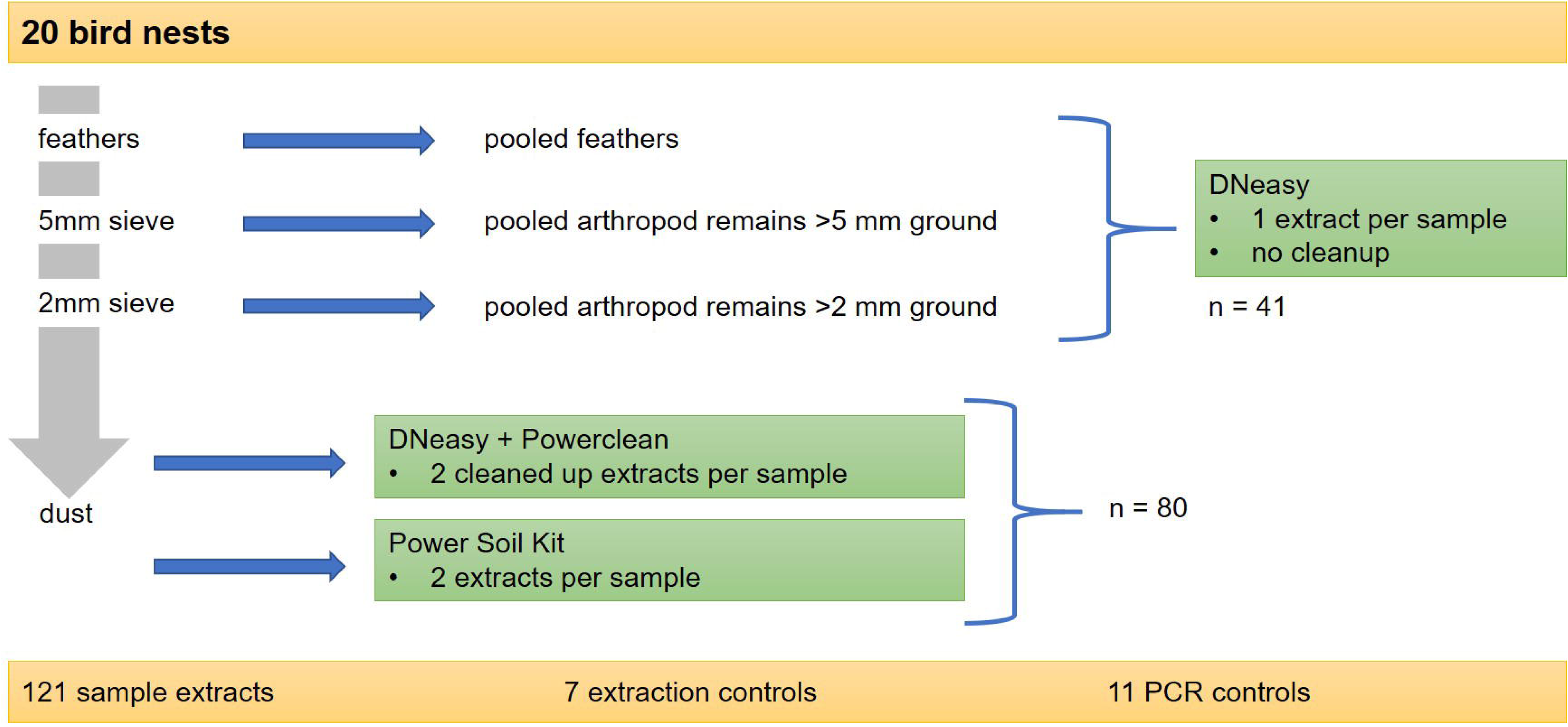
Workflow describing the processing of the bird nests for molecular analyses

### Lysis and extraction

Arthropod remains were ground to fine powder. For larger samples we used 40 ml grinding chambers and an IKA Tube Mill control (IKA, Breisgau, Germany) at 25,000 rpm for 2 × 3 min. Smaller samples were ground in 20 ml homogenization chambers using the IKA ULTRA-TURRAX Tube Drive Control System (IKA, Staufen im Breisgau, Germany) at 4,000 rpm for 30 min with 10 steel beads (diameter, 5 mm). All samples were covered with a 20:1 mixture of TES (0.1 M TRIS, 10mM EDTA, 2% sodium dodecyl sulphate; pH 8) and Proteinase K (Qiagen, 20 mg/mL), thoroughly vortexed, and incubated at 56°C overnight. Subsequently, we used 200 µl of lysate per sample for DNA extraction with the DNeasy kit (Qiagen) following manufacturer’s instructions, except for the elution step, which was done twice using 50 µl AE buffer per sample. Feather samples were lysed with 300 µl of lysis buffer (20 : 1 ratio for TES and Proteinase K). Incubation and DNA extraction were identical to the ground arthropod samples.

We generated four (and in exceptions 5) DNA extracts for each nest dust sample: two with the DNeasy PowerSoil Kit (Qiagen) and two with the DNeasy kit followed by processing with the DNeasy PowerClean Pro Cleanup Kit (Qiagen). For four dust samples, three PowerSoil extractions were carried out. First, 2 × 0.25 g dust were processed individually using the DNeasy PowerSoil Kit. Instead of the 10 min vortex step for cell lysis as suggested in the manufacturer instructions, samples were vortexed only briefly and then incubated at 70°C for 5 min. This process was carried out twice before returning to the remainder of the standard protocol. The remaining nest dust was separated into two 50 ml falcon tubes (in case of large dust samples the amount was limited to 10 g of dust per tube) and covered with a 20:1 mixture of TES and Proteinase K. The lysis and extraction with the DNeasy kit were the same as for the ground arthropod samples, but the generated extracts were additionally subjected to inhibitor removal with the DNeasy PowerClean Pro Cleanup Kit (Fig. 2). For each sample type and extraction, at least one negative extraction control was processed along the samples. Success of the extractions was confirmed by measuring the total DNA concentration with the Qubit dsDNA HS Assay Kit (Thermo Fisher Scientific).

### Amplification, library preparation, and sequencing

All generated DNA extracts were subjected to a two-step PCR protocol (Elbrecht & Steinke, 2018) with total PCR volumes of 25 µl using the Multiplex PCR Master Mix Plus (Qiagen), 0.5 mM of each primer, and molecular grade water. The primers BF3 (5’ CCHGAYATRGCHTTYCCHCG 3’) and BR2 (5’ TCDGGRTGNCCRAARAAYCA 3’) and 4 µl of DNA extract were used for the amplification of the cytochrome oxidase I (COI) gene in the first PCR (Elbrecht et al., 2019; Elbrecht & Leese, 2017). The second PCR was carried out using fusion primers containing sample-specific inline tags and 2 µl of PCR product generated in the first PCR (Elbrecht *et al*. 2019; Elbrecht & Leese 2017; Elbrecht & Steinke 2019). The thermocycling conditions for both PCRs were 95°C for 5 minutes, 35 (1^st^ PCR) or 24 (2^nd^ PCR) cycles of 95°C for 30 seconds, 50°C for 30 seconds, and 72°C for 50 seconds with a final extension at 68°C for 10 minutes. A total of 11 PCR negative controls were used and after PCR 2 the amplification success was checked on 1.5% agarose gels.

For amplicon normalization we processed 20 µl of each PCR 2 product using the SequalPrep Normalization Plate Kit (Invitrogen). We pooled 5 µl per normalized sample into a 5 ml reaction tube. After pooling, its contents were vortexed and distributed into three 1.5 ml reaction tubes. These went through clean up using the SPRIselect Kit (Beckman Coulter) and the Left Side Size Selection procedure with a sample-to-volume ratio of 0.75 and final elution in 30 µl molecular grade water. The eluates of the three reaction tubes were pooled again, the resulting library was checked on an 1.5% agarose gel and its DNA quantity was measured five times with the Qubit dsDNA HS Assay Kit (7.1 ng/µl average). Sequencing was carried out at the University of Guelph’s Advanced Analysis Centre using an Illumina MiSeq with the 600 cycle Reagent Kit v3 (2 × 300) and 5% PhiX spike in. Sequencing results were uploaded to the NCBI Sequence Read Archive (SRA, Genbank, accession: SRR23716567)

### Sequence processing

Obtained sequences were processed with JAMP (https://github.com/VascoElbrecht/JAMP). During demultiplexing, only sequences with perfectly matching tags were considered for further processing. Forward and reverse reads were merged with U_merge_PE() accessing Usearch (Edgar 2010) requiring 75% of the bases to match. After removing the primer sequences with Cutadapt (default settings; Martin 2011) and a preliminary check of the read length distribution in the remaining dataset, fragments with lengths between 200 and 500 bp were selected for further processing. An expected error value of 2 was used to remove sequences with poor quality as implemented in Usearch (Edgar & Flyvbjerg 2015). During the denoising process, reads were dereplicated per sample and amplicon sequence variants (ASV) with less than 5 reads per sample were removed, as were ASV below 0.001% relative abundance in at least one sample. The obtained ASV were mapped against a custom sequence database consisting of all publicly available COI sequences on BOLD (as of May 5^th^ 2021) assigned to “Ontario” using the USEARCH algorithm. All reference sequences were degapped and ambiguous base calls of more than seven Ns in a row were removed prior to mapping. Our primary aim was to identify Annelida, Arthropoda, Chordata, and Mollusca. As the BOLD COI barcode database is very comprehensive for these taxa in southern Ontario, we refrained from separately assigning sequences to different taxonomic levels based on similarity percentages. Only ASV with at least 99% similarity were used and assigned to species level (Elbrecht *et al*. 2017b). All matched ASV were subjected to a manual plausibility check. This resulted in the use of genus-level information if a) COI was not suitable to distinguish between individual species or b) if the detected species does not occur in the study area, but the genus does. ASV for which no species and/or genus name were available, but which passed this similarity threshold during mapping, were kept in the dataset. For any sequences detected in negative controls, twice the number of reads were removed from all co-extracted samples containing the respective species.

### Reason for detection

Each of the detected taxa was assigned to a category of reasons for occurring in a bird nest: “bird food”, “feeding in nest”, “contained in nest material”, “nest material”, “parasite” and “visitor”. Additionally, each taxon was associated with a consumer type based on the behavior at its adult life stage: “decomposer”, “not eating”, “palynivore”, “fungivore”, “herbivore”, “omnivore”, “parasite”, “predator”, “saprophagous” and “scavenger”. For both categorisations, an unambiguous assignment was not always possible.

### Statistical analysis

All calculations and visualizations were made in R Version 4.2.2 (R Core Team 2021) using the packages “ggplot2” (Wickham 2016), “dplyr” (Wickham *et al*. 2022), “ggrepel” (Slowikowski 2021) and “ggpubr” (Kassambara 2020). Poisson regressions with a log-link function were used to test for significant differences in the number of detected taxa between the four different extract types (ground arthropods, dust DNeasy extracted, dust Power Soil extracted, feathers). The unbalanced distribution of sampled nests between bird species did not enable the application of ordination methods to examine differences between species communities with respect to the nesting bird and resulted in the use of descriptive statistics for the obtained data.

## Results

During the consecutive sieving and sorting process of the 20 nests, 32 samples of ground arthropods were generated, 14 of which contained remains larger than 5 mm. Nine nests contained feathers and for each nest, four dust samples were processed (two per extraction method; Fig. 2).

### Sequence processing and taxonomic assignment

The Miseq run produced 27,407,037 reads and 9.69% of these were discarded during demultiplexing. Of the discarded sequences, 19.4% were assigned to PhiX using a similarity threshold of 90%. On average, 76.5% of the forward and reverse reads per sample could be merged and the removal of the primer sequences was successful for 99% of the reads. After length selection, 2,523,299 reads, on average 14.5% per sample, remained for further processing. During the denoising 1,393,893 reads were removed (including chimeras) and 791,123 reads were used for taxonomic assignment. The extraction controls processed alongside the ground arthropods and the feather samples showed contaminations with *Turdus migratoris* Linnaeus, *Lumbricus terrestris* Linnaeus, *Lymantria dispar* (Linnaeus), and *Ornithonyssus sylviarum* (Canestrini and Fanzago). After down-correcting the reads in the affected samples, removing *Homo sapiens* sequences, and a general plausibility check, 352,692 reads belonging to 103 taxa were used for the analysis.

Ground arthropod samples and Power Soil extractions each accounted for 39% of the reads, whilst DNeasy extractions and feather samples accounted for 13% and 9%, respectively. Aves (68%) and Insecta (23%) were the two classes to which most reads were assigned to (Fig. 3).

**Figure 3.**
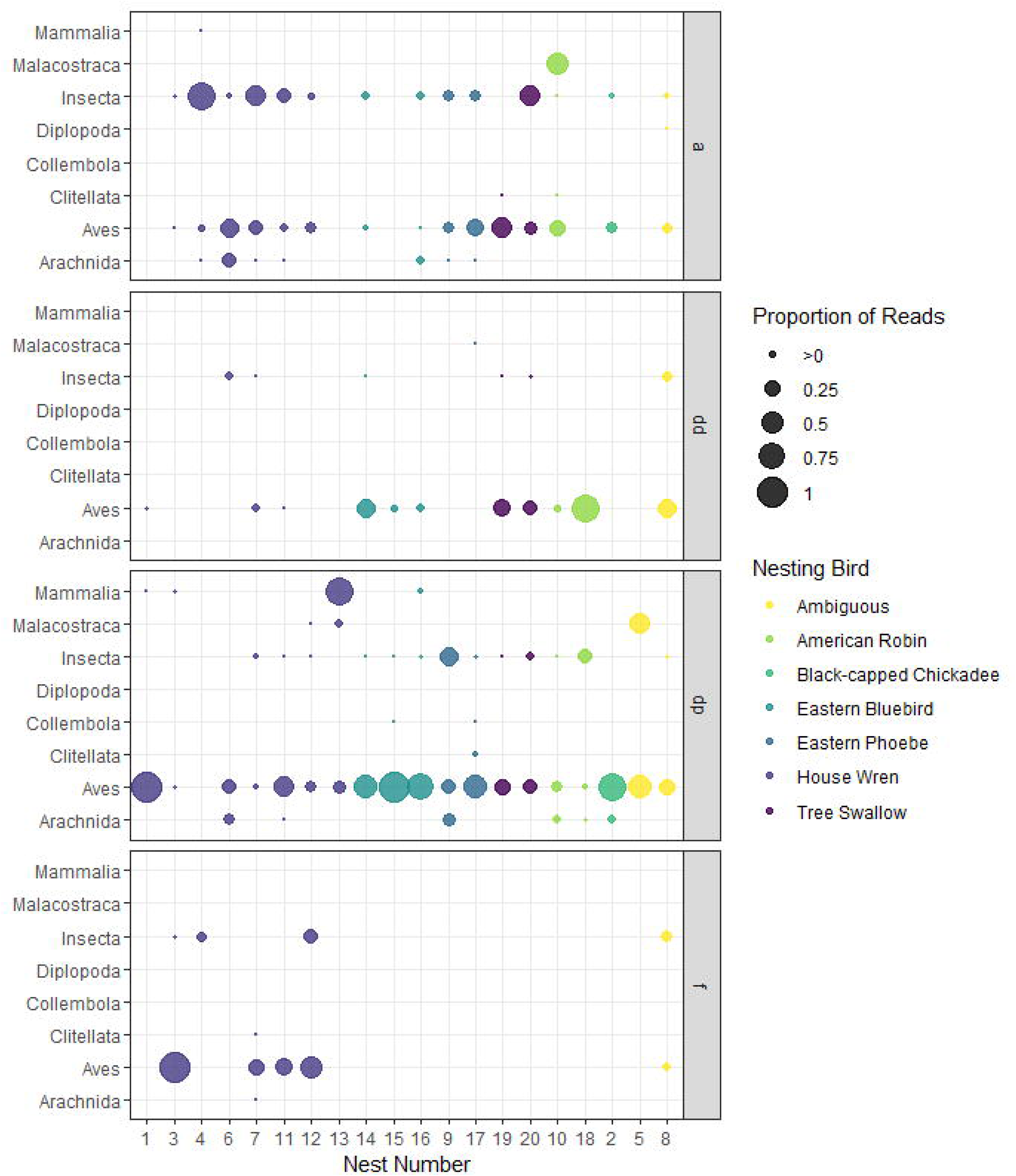
The bubble plot shows the proportion of reads per bird nest which are associated to the four different sample/extraction types (“a”: ground arthropods, “dd”: DNeasy extraction, “dp”: Power Soil extraction, “f”: feather samples), i.e., proportions for each nest sum up to 1. Nests are sorted and bubbles are coloured according to the nesting bird species and the proportional reads are separately displayed for all detected taxonomic classes.

### DNA-based detections in subsamples and sample types

For nine nests, taxonomic assignments were possible for small and medium ground arthropod subsamples. Of the taxa detected in these samples (mean 9.11 ± 5.86 SD), an average of 5.5% were simultaneously detected in both subsamples. Both subsamples of the DNeasy extraction led to taxonomic assignments for 10 nests, with a mean of 3.10 ± 2.02 SD taxa of which on average 75.50% were detected in both subsamples. The Power Soil extraction enabled taxonomic identifications in more than one subsample for 17 nests. Of the detected taxa (mean: 4.82 ± 2.24 SD), 47.31% were on average detected in more than one subsample per nest.

Of 19 nests with taxonomic assignments based on the Power Soil extraction, 13 had DNeasy-based detections and 62.86% of these (DNeasy detections) were observed with both methods. Per nest, significantly more taxa were detected from dust with the Power Soil extraction method compared to the DNeasy extraction method (Table 2; mean Power Soil: 4.89 ± 2.28 SD; mean DNeasy: 2.77 ± 2.05 SD). A comparison of the number of detected taxa per nest and sample type revealed significantly less detected taxa in DNeasy extractions compared to the other three sample types (Table 2; Fig. 4).

**Table 2.** The Generalized Linear Model (Poisson error distribution, log-link) describing the relation between the number of detected taxa in the four sample types: ground arthropods, DNeasy extracted nest dust and Power Soil extracted nest dust samples, and feather samples. DNeasy extractions were used as the base category.

|  | Estimate | Std. Error | z-value | p-value |
| --- | --- | --- | --- | --- |
| (Intercept) | 0.99 | 0.17 | 5.86 | <0.001 |
| ground arthropods | 1.01 | 0.19 | 5.22 | <0.001 |
| Power Soil extractions | 0.60 | 0.20 | 3.01 | <0.01 |
| feathers | 0.86 | 0.23 | 3.65 | <0.001 |

**Figure 4.**
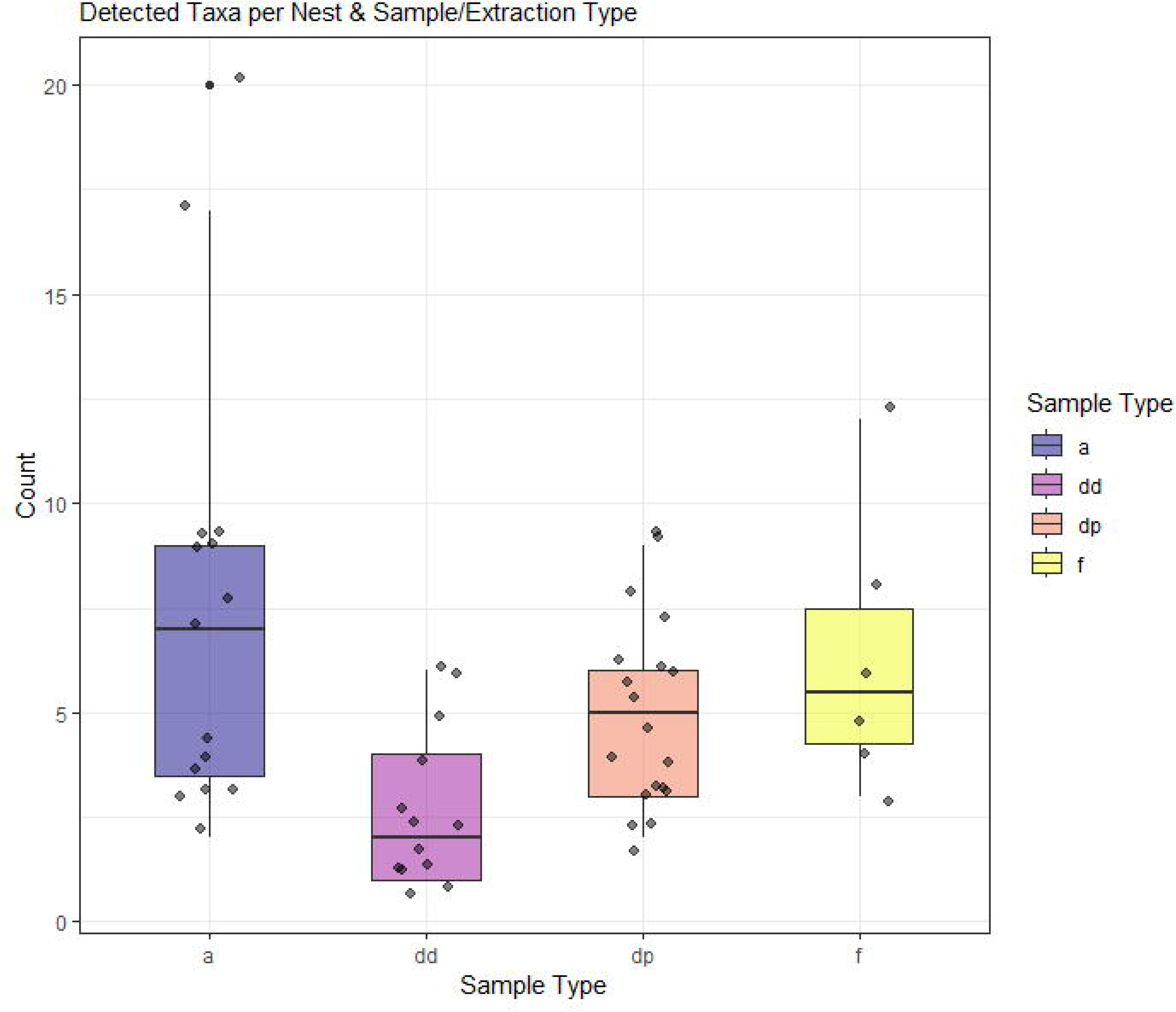
Box plots of the detected taxa in the four sample types: ground arthropods (a), DNeasy extracted nest dust (dd), Power Soil extracted nest dust samples (dp), and feather samples (f). Significantly less taxa were detected after DNeasy extraction of dust (Table 2).

### Detected biodiversity

In the four nests processed using emergence traps, 24 distinct taxa were detected of which 58% were identifiable to species level using DNA barcoding. Insecta made up 92% of the detected taxonomic diversity in emergence traps with Diptera, Coleoptera and Psocodea each contributing 25%, 25%, and 21%, respectively (Fig. 5). The metabarcoding analysis led to the detection of 103 distinct taxa, 84 of which were identifiable to the species level (See Supporting Information 1 for the full data table). Arthropoda and Insecta accounted for 72% and 54% of the distinct taxa detected on the phylum-level and class-level, respectively. At the order-level, Diptera and Passeriformes accounted for the highest percentage of taxonomic diversity (18% and 14%, respectively, Fig. 5). Five taxa (*Niditinea* Petersen, *Ceratophyllus* Curtis, *Protocalliphora* Hough*, Gaurax pallidipes* Malloch, and *Liposcelis corrodens* (Heymons)) were detected both in emergence traps and with DNA metabarcoding.

**Figure 5.**
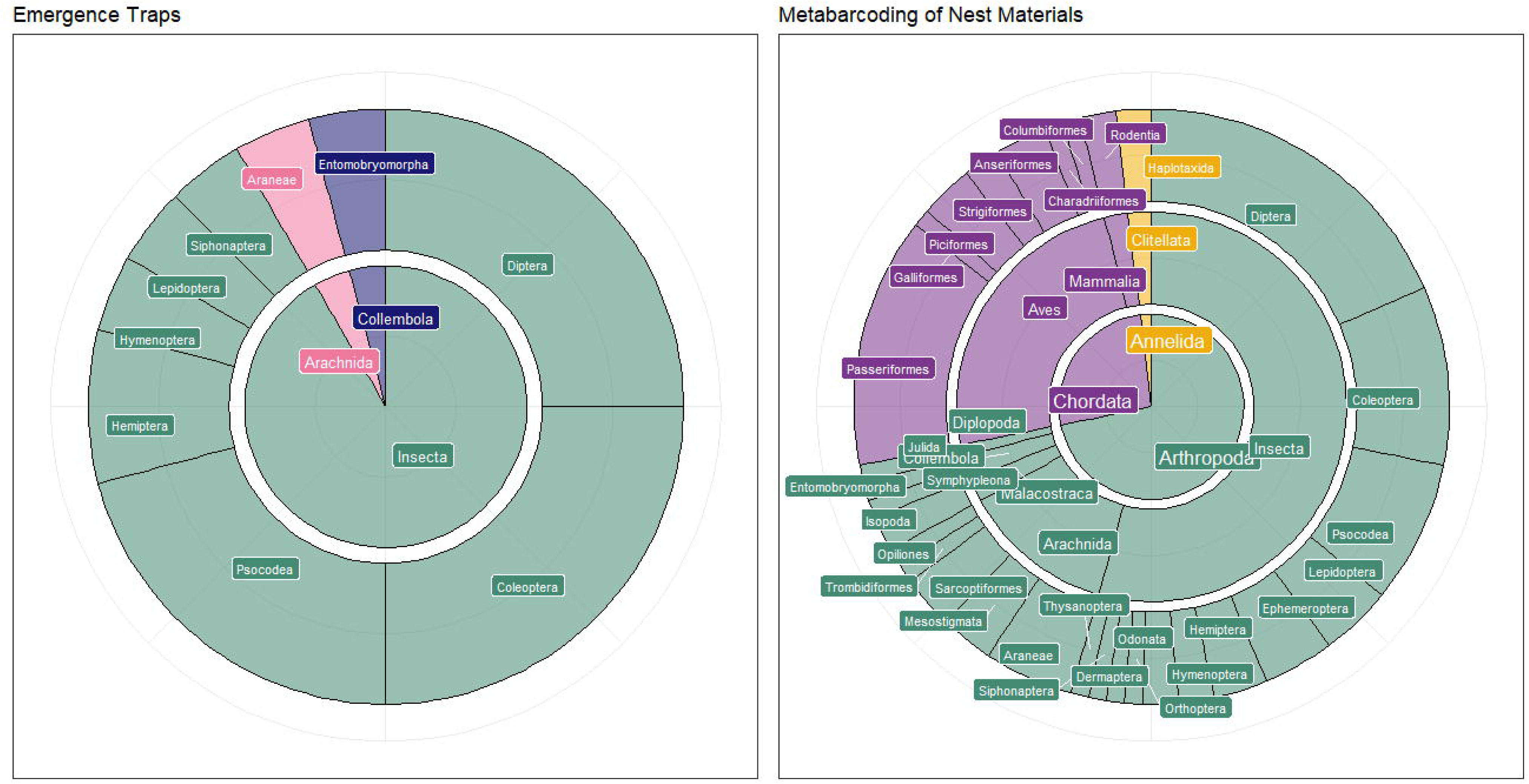
Taxonomic diversity (i.e., proportion of distinct taxa) detected for the emergence trap’s arthropods and the metabarcoding of nest materials. Emergence trap data were obtained from four nests; metabarcoding data from 20 nests and 121 extracts.

Altogether, 25 bird species, all of which occur in the study region, were detected with metabarcoding. Except for the Black-Capped Chickadee (*Poecile atricapillus* (Linnaeus)), the nesting bird species was always detected in the respective nest(s) and accounted for 32-71% of the total Aves reads (summed up per nesting species; Fig. 6). Most bird species (n=18) were detected in the eight House Wren (*Troglodytes aedon* Vieillot) nests.

**Figure 6.**
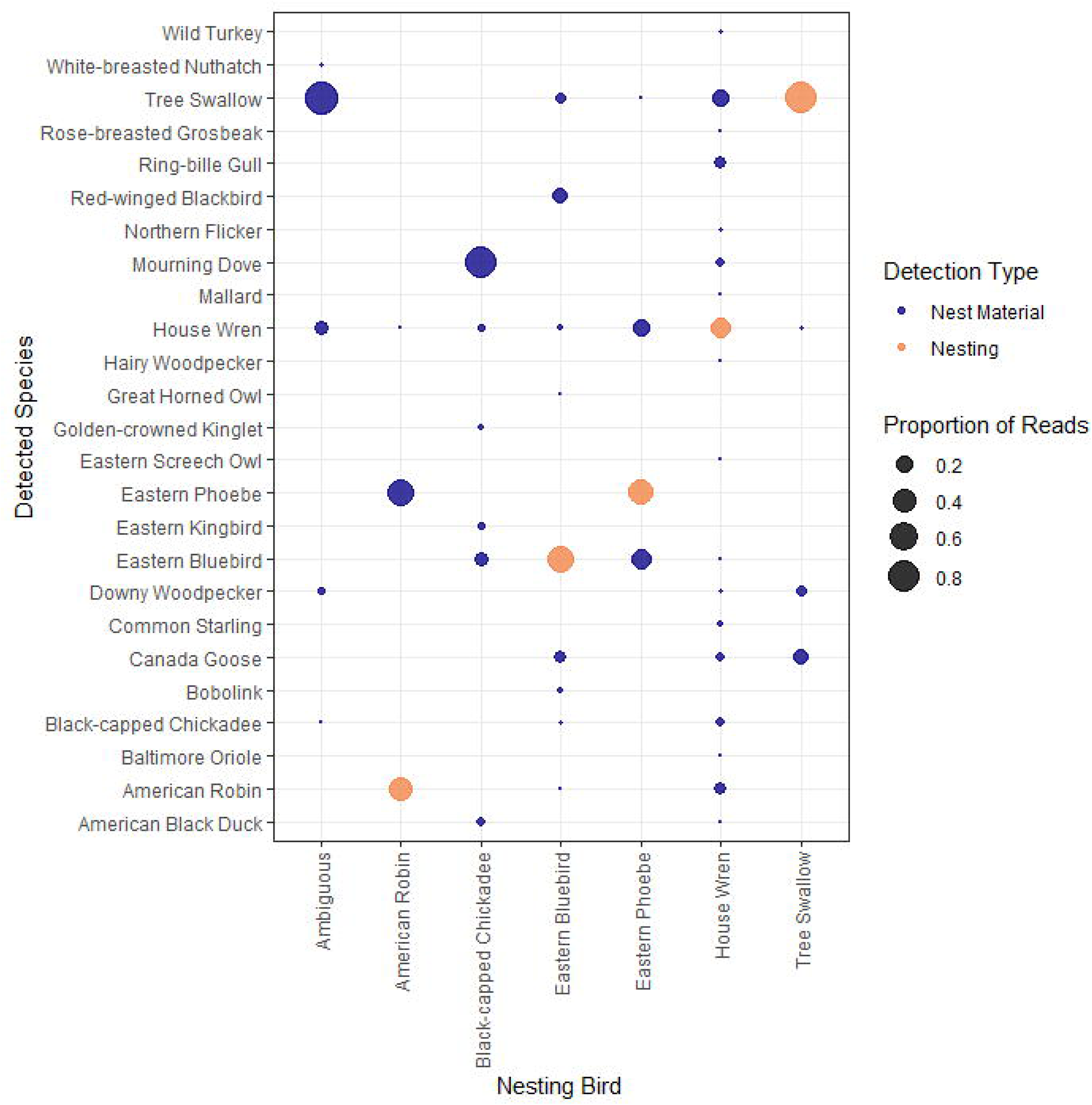
The bubble plot shows the proportion of reads which are assigned to one of the 25 detected bird species for every nesting bird species i.e. proportions for each nesting bird sum up to 1. Detections of the nesting bird species itself are coloured in orange, detections of species which are foreign to the nest, are coloured in blue.

### Functional role of detected taxa

Of the 103 taxa detected with DNA metabarcoding, 73% could be associated with a specific reason for occurring in the nest samples: the occurring Arthropoda were primarily food for the nesting birds or feeding in the bird nest whilst the detected Chordata (i.e. Aves and Mammalia) were categorized as part of the nest material or as visitors (Fig. 7). With respect to consumer type, 68% of the detected taxa had an unambiguous link at the adult stage: the detected Annelida were identified as decomposers, whilst most of the detected Arthropod taxa could be associated with herbivory (Fig. 7).

**Figure 7.**
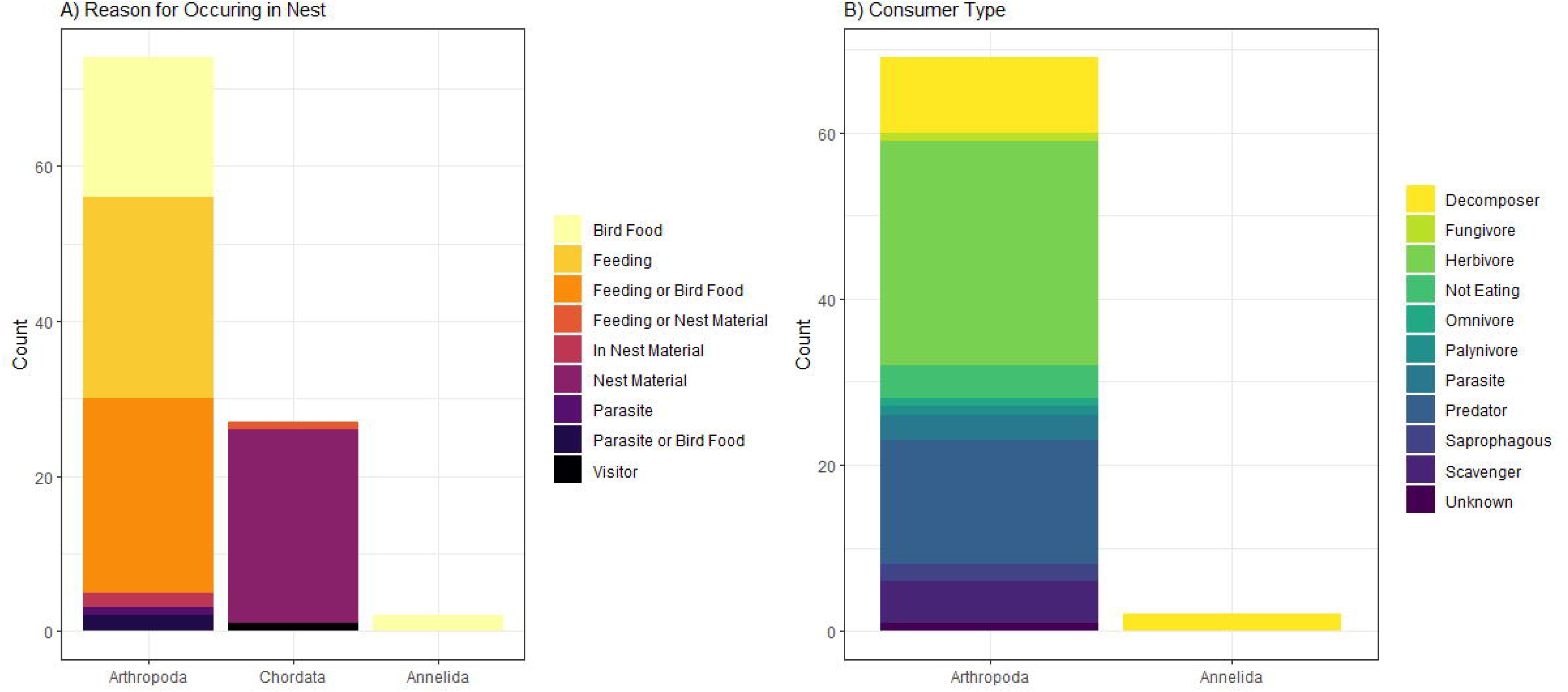
Bar plot showing the number of taxa per phylum which could be unambiguously assigned to A) a reason for occurring in a bird nest (n=75) and B) a consumer type (n=70).

## Discussion

This work demonstrates the potential of molecular analysis of animal remains and eDNA contained in bird nests. Our metabarcoding approach delivered a detailed picture of the taxonomic diversity accumulated in bird nests. In comparison to data from four emergence traps, metabarcoding of nest dust, arthropod remains and feathers from 20 nests detected more than four times as many taxa. Overall, ground arthropod samples and nest dust analyzed with the PowerSoil kit were most informative, resulting in an average detection of 5.90 and 6.25 taxa per nest, respectively. Detections from nest dust eDNA were possible for all but one nest and the number of detected taxa varied less across the investigated bird species in comparison to results obtained from arthropod remains. Of the two extraction protocols tested with nest dust eDNA, extracts processed with the PowerSoil kit detected significantly higher numbers of taxa per nest than the DNeasy kit in combination with the PowerClean Pro inhibitor removal. Nevertheless, detections in subsamples from the same nest were more homogenous with the latter approach. Ultimately, it was possible to obtain data for a wide range of taxa constituting bird food, parasites, nest materials, and foraging in the nests.

The taxonomic diversity varied greatly between the emergence traps and metabarcoding data with only five taxa overlapping between both methods and eight joint detections of a taxon in both the emergence trap and the metabarcoding data of the same nest (Fig. 5). The emergence trap only targets alive arthropods in the nests at the point of collection (Levesque-Beaudin et al., 2020), while the eDNA-based approach can capture alive and dead organisms as well as fragments of organisms (Barnes & Turner, 2016). The eDNA-based data thus represents a larger time frame, provided the DNA traces withstand environmental conditions (Oehm et al., 2011). Additionally, the long activation period of the emergence traps (two weeks at room temperature) might have induced further eDNA decomposition.

Overall, the highest number of taxa was detected from ground arthropod remains (n=67) and nest dust analyzed with the PowerSoil kit (n=23), albeit the majority of reads were attributed to different groups: for ground arthropod remains, most reads were assigned to Arthropoda; for dust samples, Aves showed the highest read numbers. The total number of detected taxa varied more for ground arthropod remains than for nest dust samples extracted with the Power Soil kit, with arthropod remains not available or not resulting in molecular detection for 5 nests contrasting with a maximum of 20 taxa per nest. These differences can be attributed to distinctions in the choice of nest materials and nest sanitation between bird species (Collias, 1997; Guigueno & Sealy, 2012; Mennerat et al., 2009). For instance, nests of House Wrens which are primarily built from twigs (McCabe, 1965), are more difficult to clean and thus potentially contain more arthropod remains than nests built from soft plant materials and clay.

Generally, ground arthropods are a great source of DNA, even old museum specimens can provide sequences (Prosser *et al*. 2016), and DNA degradation has a limited effect on community recovery (Krehenwinkel *et al*. 2018). To avoid large specimens contributing an excessive amount of DNA (Elbrecht *et al*. 2021a; Krehenwinkel *et al*. 2018) and to maximize the chances of detecting rare taxa, it is, however, advisable to process them in different size categories to increase recovery (Creedy *et al*. 2019; Elbrecht *et al*. 2017a, 2021a). Other studies have used 3 to 6 size categories (Elbrecht *et al*. 2017a, 2021a; Krehenwinkel *et al*. 2018). In the present study, two size categories were deemed sufficient to reduce sorting effort, laboratory time and costs. Albeit it is not possible with the present dataset to determine any positive effects of the size sorting, with only 5.5% of the taxa being detected in both size categories indicates a clear taxonomic distinction between the two groups.

Feathers are an excellent source of bird DNA (McInnes *et al*. 2021; Smith *et al*. 2003), but also of others animals associated with them such as feathers mites (Diaz-Real *et al*. 2015; Doña *et al*. 2019a; b). They not only provide information on the nest inhabitants, but also visitors, as well as the local bird diversity through feathers selected for nest building (Collias & Collias 1984; Healy *et al*. 2015). Of the 20 examined nests, nine contained feathers and six of these samples resulted in DNA-based detections of Aves, Insecta and Clitellata (3-12 taxa per sample). Despite these promising results, feather sampling alone might not be sufficient for future studies, since not all bird species use feathers as nest materials and the majority of arthropod taxa detected from nest dust were not detectable in feather samples.

Dust is known to be a good source of plant eDNA (Lennartz *et al*. 2021), but can also be used to detect arthropods (Krehenwinkel *et al*. 2022) and vertebrates (Lynggaard *et al*. 2022). In comparison to metabarcoding results obtained from bulk samples of terrestrial or aquatic arthropods (Gleason et al., 2021; Steinke et al., 2021), the number of high-quality reads obtained from the different sample types in this study, is low. Only 1.3% of all reads were ultimately used for taxonomic assignment. Provided that we employed a combination of primers and bioinformatic processing, which is well established for arthropod metabarcoding, there are three main reasons for this turnout: i) the DNA of Arthropoda and Chordata contained in dust samples is difficult to amplify because it’s likely only a fraction of the plant and microorganism DNA contained in theses samples. Under such conditions, selecting a primer pair which is less prone to amplify plant, fungal, and microbial DNA is crucial (Elbrecht *et al.,* 2019, Krehenwinkel *et al*. 2022). ii) Much of the target DNA in the dust and in arthropod remains is likely old and degraded; it gradually accumulated during the breeding season and was exposed to decay processes. Generally, DNA degrades quickly with UV light and rain (Oehm et al., 2011), but dry conditions in nest boxes likely slow the degradation process (Matange *et al*. 2021) and thus enable the detection of DNA traces accumulated over a longer time period. iii) The abundance of plant materials, clay and traces of bird feces potentially made the samples prone to inhibition (Sidstedt et al., 2020; Thalinger et al., 2017). Both the PowerSoil kit and the DNeasy kit in combination with the PowerClean inhibitor removal kit resulted in successful amplifications in PCR from nest dust eDNA samples. DNA extracts obtained with the PowerSoil kit detected significantly more taxa than extracts processed with the DNeasy and PowerClean kits, but detections were more homogenous between subsamples of the latter. This likely results from the creation of one large lysate prior to DNeasy extraction, which improved mixing of the DNA. In contrast, the PowerSoil kit processes only a small amount of dry material and taxonomic overlap between subsamples was considerably smaller (DNeasy: 75.5% vs. PowerSoil: 47.31%). To conclude, we suggest processing several subsamples per nest if only small amounts of dust are extracted. Despite these limitations, the diversity of taxa detected from nest dust exceeded detections obtained from the emergence traps and feathers. For nest types which hardly yield arthropod remains at the end of the breeding season, we consider the analysis of eDNA from nest dust a viable alternative to processing with emergence traps.

Most arthropods detected in the nest were either predators or prey (Fig. 7). As most nests came from insectivorous bird species, it is not surprising that the majority serve as bird food. Regarding the associated trophic roles, the dataset was dominated by herbivores, predators and decomposers. All these create a chain of interactions within the same nest: herbivores feeding on nest materials, predators feeding on herbivores, and decomposers cleaning up the leftovers. Hence, sampling a bird nest provides a good portrait of the nest micro-ecosystem and its surrounding area. The metabarcoding approach also provided insight into the bird communities by identifying feathers that were used as nesting material, left behind by possible nest visitors or resulted from specific bird species behavior. Eastern Bluebirds (*Sialia sialis* (Linnaeus)) like to line their nest with fine grasses, hairs or feathers (Cornell Lab of Ornithology 2019a), hence the detection of eight additional bird species was not surprising (Fig. 6). The American Robin nest had a large proportion of reads matching Eastern Phoebe. These birds like to reuse nests, even those of Robins (Cornell Lab of Ornithology 2019b). It is possible that a Phoebe visited the nest or had used it previously. The large number of species detected in the House Wren nests (n=17) potentially stem from their nest-destroying behaviour (Belles-Isles & Picman 1986; Pribil & Picman 1991). As a result, they return with DNA traces or fragments of the nest materials to their own nest.

## Conclusion

Metabarcoding of arthropod remains and eDNA in dust samples is a viable alternative to studying bird nest communities solely via emergence traps. These samples can capture a much broader taxonomic range and are not restricted to live organisms. Additionally, the metabarcoding approach allowed for the identification of fur and feathers used as nest materials, remains of birds’ dietary samples, species living exclusively in bird nests, and bird parasites. Arthropod remains and nest dust eDNA extracted with the PowerSoil kit were the most promising approaches for upscaling the assessment of bird nest communities in the future to systematically investigate functional diversity in nests and differences between bird species.

## Acknowledgments

We thank Chris Earley for providing access to the nest boxes of the University of Guelph Arboretum.

## Data availability

The raw metabarcoding data has been uploaded to GenBank SRA (accession: SRR23716567). All data obtained from emergence traps and molecular data used for the consecutive analyses have been uploaded to Figshare and are available at https://doi.org/10.6084/m9.figshare.22203616.v1

## Author contribution statement

VLB conceived the original idea, which was developed further together with DS and BT. DS secured the necessary funding. VLB collected the bird nests and was responsible for the emergence traps and the consecutive morphological identification of emerging arthropods. VLB and BT sieved the nests and sorted the arthropod remains, BT was responsible for the consecutive laboratory processing of the generated samples which was carried out together with MB. BT was responsible for analyzing the generated data, VLB wrote a first draft of the manuscript to which all co-authors contributed critically before final approval for publication.

